# *beditor*: A computational workflow for designing libraries of guide RNAs for CRISPR-mediated base editing

**DOI:** 10.1101/426973

**Authors:** Rohan Dandage, Philippe C Després, Nozomu Yachie, Christian R Landry

## Abstract

CRISPR-mediated base editors have opened unique avenues for scar-free genome-wide mutagenesis. Here, we describe a comprehensive computational workflow called *beditor* that can be broadly adapted for designing guide RNA libraries with a range of CRISPR-mediated base editors, PAM recognition sequences and genomes of many species. Additionally, in order to assist users in selecting the best sets of guide RNAs for their experiments, *a priori* estimates, called *beditor* scores are calculated. These *beditor* scores are intended to select guide RNAs that conform to requirements for optimal base editing: the editable base falls within maximum activity window of the CRISPR-mediated base editor and produces non-confounding mutational effects with minimal predicted off-target effects. We demonstrate the utility of the software by designing guide RNAs for base-editing to create or remove thousands of clinically important human disease mutations.

## INTRODUCTION

CRISPR-mediated base editors (BEs) are engineered by fusing a DNA modifying protein with a nuclease-defective Cas9 (dCas9) protein, allowing scar-free targeted mutagenesis (1–5). Currently, two major types of BEs are available – cytosine base editors (CBEs) that enable the conversion of cytosine to uracil by catalysis and then to thymine through replication or repair, (1, 6–8)–for example BE3 (6) and Target-AID (C•G to T•A) (1). Similarly adenine base editors (ABEs) enable conversion of adenine to inosine by catalysis and then to guanine through replication or repair (A•T to G•C) (2, 9, 10)–for example ABE7.10 (2). Currently BEs enable many codon level substitutions and thus amino acid substitutions (Figure S1). Owing to the unique capability of scar-free mutagenesis, BEs have recently found numerous applications in both model and non-model organisms (3, 10–13) and have substantial promise in therapeutic applications (14, 15).

For designing guide RNAs (gRNA) in BE mediated mutagenesis experiments, several specific requirements for the optimal activities of DNA modifying and Protospacer Adjacent Motif (PAM) sequences need to be taken into consideration. The complexity of this task especially increases when designing gRNA libraries against large sets of targets located across a given genome. Because design of gRNA sequences is a primary requirement for the success of the BE-mediated mutagenesis experiment, the methods and strategies involved in designing gRNA libraries are arguably the one of the most important factors in such experiments.

Currently available gRNA designing tools are however either specifically focused on non-sense mutations (16) or allow very limited customization (BioRxiv: https://doi.org/10.1101/373944) (See Table S1), leaving a major roadblock in the applications of the BEs in genome editing. The continuous discoveries of new BEs and the expansion of their existing editing capabilities (17) demand a complementary development in computational methods that would allow designing gRNAs using new and improved base editors. Moreover, considering the prospective applications across non-model organisms, compatibility with diverse genomes is essential. Overall, therefore, a comprehensive computational framework to design gRNAs libraries can potentially fuel the progress in CRISPR-base editing technology and its diverse applications in genome editing.

We developed a comprehensive computational workflow called *beditor* (Figure 1a) that can design gRNA libraries with any requirements of DNA modifying enzyme. These include the range of nucleotides where maximum catalytic activity of BE occurs, henceforth simply referred to as ‘activity window’, and PAM recognition sequence. *beditor* is directly compatible with more than 125 genomes hosted in the Ensembl genome database (18) and any annotated custom genomes. Additionally, the *beditor* workflow also provides *a priori* estimates called *beditor* scores for each gRNA that accounts for specific editing requirements of BEs, gRNA binding at off-target sites and the number and types of off-target effects (Figure 1b). Such estimates will inform researchers in optimizing throughput of their BE-mediated mutagenesis experiments. With its Graphical User Interface (GUI), command line interface and open source Application programming interface (API), the *beditor* workflow has broad applicability for the genome editing community and its open source implementation will allow for continuous enhancements in the future.

**Figure 1.**
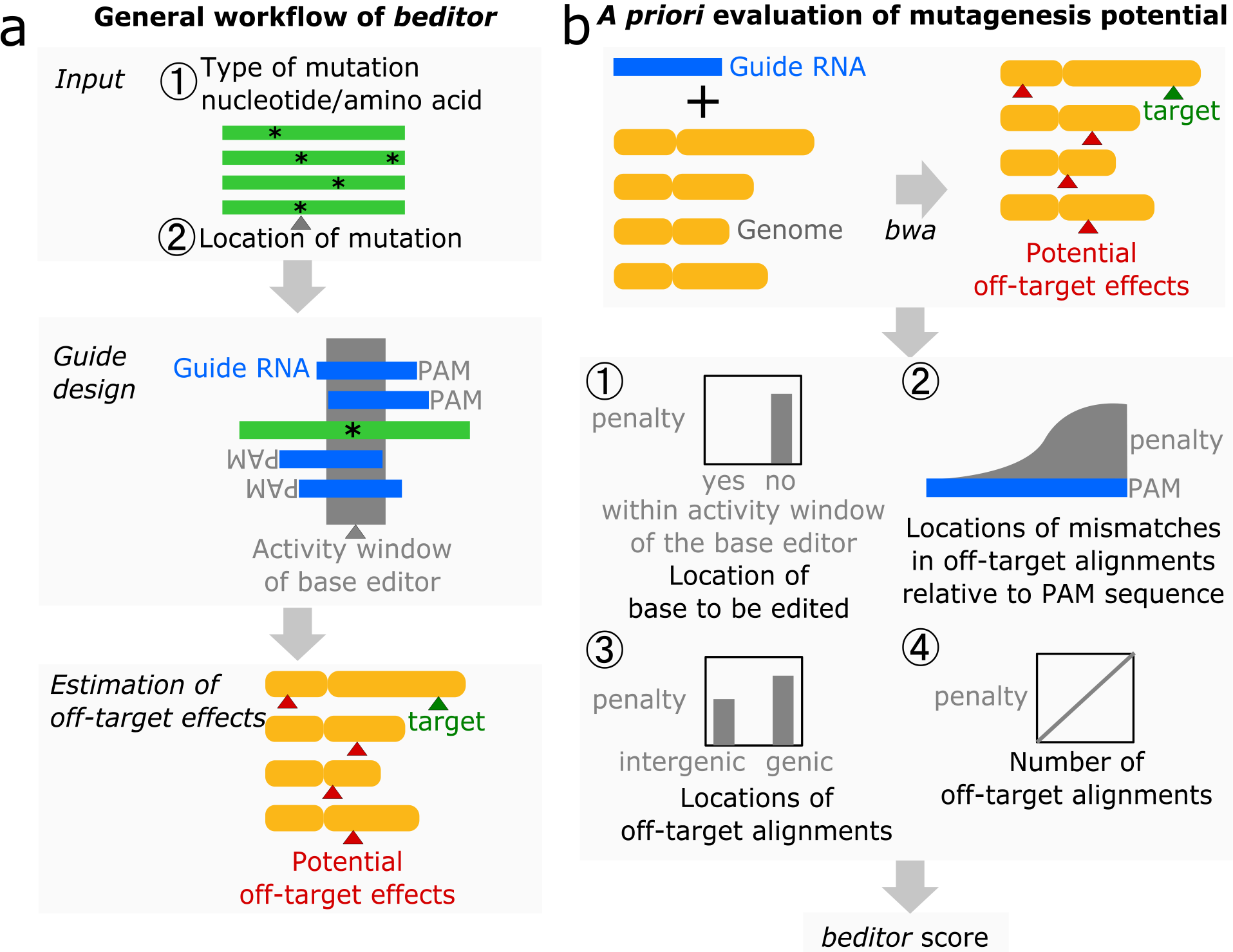
The computational workflow of *beditor* allows for the flexible design of gRNA libraries to be used in CRISPR base editing and offers *a priori* evaluation of mutagenesis potential. **a** Information on the type and location of desired mutations is supplied to the *beditor* workflow as a tab separated file. gRNAs are designed according to the user provided sets of BEs and PAM recognition sequences. Among many base editor and Pam sequence specific requirements, nucleotide windows for maximum activity are considered while designing the gRNAs. Finally, potential off-target effects are estimated. **b** A scoring system specifically designed for *a priori* evaluation of mutagenesis potential of gRNAs. Penalties are assigned based on (1) the total number of off-target alignments of gRNAs to the reference genome, (2) positions of the mismatches in the off-target alignments relative to the PAM and (3) genomic locations of off-target alignments and lastly, (4) whether the editable base lies inside the activity window of the BE. Using all of the above penalties, a final score is calculated for each gRNA sequence – the *beditor* score.

## MATERIALS AND METHODS

### Implementation

The *beditor* workflow is implemented as an open-source python 3.6 package hosted at https://pypi.org/project/beditor. The source code of *beditor* can be accessed at https://www.github.com/rraadd88/beditor. The documentation of the software with API is available at https://www.github.com/rraadd88/beditor/README.md. The *beditor* workflow depends on other open source softwares such as PyEnsembl (19), BEDTools(20), BWA(21) and SAMtools(22) at various steps of the analysis. User provided mutation information is first checked for validity with PyEnsembl (https://github.com/openvax/pyensembl.). Genomic sequences flanking the mutation sites are fetched using BEDTools (20). The designed gRNAs are aligned the reference genome using BWA (21) and alignments are processed using SAMtools (22) for evaluation of off-target effects using the *beditor* scoring system. Visualization of alignments of guide RNAs with genomic DNA are created using the DnaFeaturesViewer package (https://github.com/Edinburgh-Genome-Foundry/DnaFeaturesViewer).

### *beditor* scoring system

Alignment of the designed gRNAs (with PAM sequence) with the provided reference genome is carried out using BWA, as used in (21), allowing for a maximum of two mismatches per alignment (23). The *beditor* score is evaluated as follows.

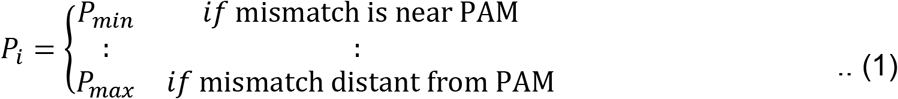

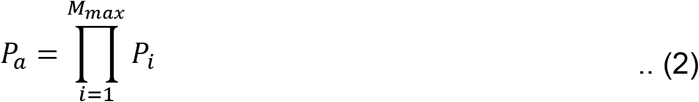

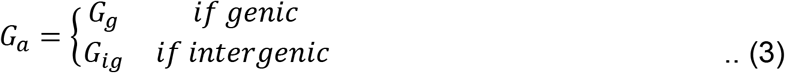

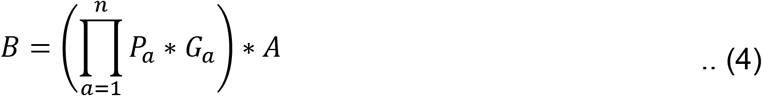

For an alignment between a gRNA sequence and the genome, *P*_*i*_ is a penalty assigned to a nucleotide in the gRNA sequence based on the position of a mismatch in the aligned sequence relative to the PAM. If the mismatch is near the PAM sequence, a minimal penalty *P*_*min*_ is assigned. Conversely, if the mismatch is far from the PAM, a maximum penalty *P*_*max*_ is assigned. The relative values of such penalties were determined by fitting a third degree polynomial equation to the mismatch tolerance data from (24) (Figure S2). This way, penalties increase non-linearly from *P*_*min*_ to *P*_*max*_, as the distance of nucleotide (*i*) from PAM sequence increases. Individual penalties assigned for all the nucleotides in a gRNA are then multiplied to estimate a penalty score for a given alignment called *P*_*a*_ (equation 2). In cases of gRNAs with lengths other than 20, the fitted equation is used to interpolate penalty scores. In case of 5’ PAMs, the order of the vector containing position wise penalty scores is reversed. *G*_*a*_ is a penalty defined by whether the off-target alignment lies within a genic or an intergenic region (equation 3). *A* is a penalty based on whether the editable base lies within the activity window of BE (equation 4).

Note that due to the lack of large scale BE editing data, in the current version of *beditor*, penalties are set based on the importance of each requirement (Table S2). Such penalties would be informed from empirical data in the future developments.

The overall *beditor* score *B* for a gRNA is determined by multiplying penalties assigned per alignment (*P*_*a*_ and *G*_*a*_) for all alignments (n) with a penalty assigned to the gRNA (A) (equation 4). Multiplication of individual penalties insures that if *any* of the criteria is suboptimal, the *beditor* score decreases.

### Demonstrative analysis 1: customizability with respect to base editing strategies

For a demonstrative analysis with custom base editors and PAM recognition sequences were used (Figure 2 and Table S3), 1000 nucleotide and amino acid mutations were randomly assigned across the genome of *S. cerevisiae* (https://github.com/rraadd88/test_beditor). Such sets of mutations create uniform datasets ideal for testing features of *beditor*. The input mutation data was created for both mutation formats (either amino acid or nucleotide) and modes of mutagenesis (“create” or “remove”). The command “beditor --cfg params.yml” was executed. Here, params.yml contains input parameters of the analysis (Table S4) in a user-friendly YAML format. For this analysis, input parameter ‘host’ was set to ‘Saccharomyces_cerevisiae’ and parameter ‘genomeassembly’ was set to ‘R64-1-1’. Summary statistics on the editability are included as Table S5 and gRNA libraries are included as Supplementary data 1.

**Figure 2.**
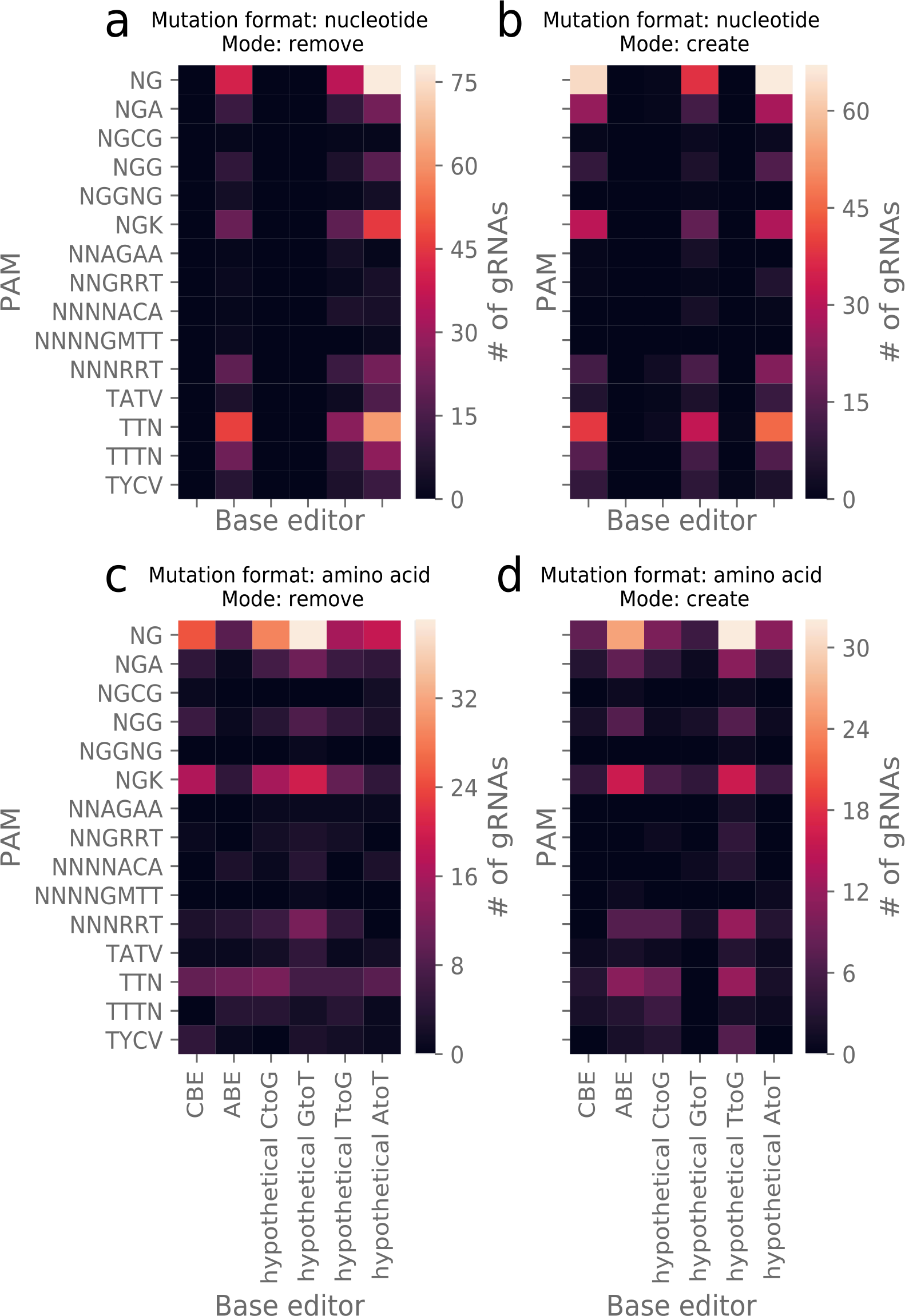
Demonstrative analysis of gRNAs designed with custom base editors and PAM recognition sequences. In order to demonstrate the utility of *beditor* in utilizing custom base editors and PAM recognition sequences, sets of 1000 randomly assigned mutations in *S. cerevisiae* (see Supplementary methods) were analyzed in 2 mutation formats (nucleotide or amino acid) and 2 modes of mutagenesis (“create” and “remove”) (each combination is shown in panel **a** to **d**). In each heatmap, number of gRNAs designed by each combination of a base editor (in columns) and PAM recognition sequence (in rows) is shown.

### Demonstrative analysis 2: ability to work with different species

For a demonstrative analysis with different species, sets of 1000 random mutations were created in genome of 10 representative species (*Bos Taurus, Danio rerio, Equus caballus, Felis catus, Gallus gallus, Macaca fascicularis, Mus musculus, Pan paniscus, Saccharomyces cerevisiae and Sus scrofa*) as described in case of demonstrative analysis 1. The input parameters used in this analysis are the same as Table S4 except for host names and genome assembly versions that were obtained from http://useast.ensembl.org/index.html. The summary statistics on the editability are included as Table S6 and gRNA libraries are included as Supplementary data 2.

### Case study analysis

For the case study analysis, a set of clinically associated human mutations were obtained from the Ensembl database in GVF format (ftp://ftp.ensembl.org/pub/release-93/variation/gvf/homo_sapiens/homo_sapiens_clinically_associated.gvf.gz, Date modified: 08/06/2018, 16:13:00). From genomic coordinates of SNPs, inputs for nucleotide mutations (reference and mutated nucleotide) and amino acid mutations (transcript ids, amino acid position, reference residue and mutated residue) were identified using PyEnsembl (19). The mutation information was provided to the *beditor* workflow as a tab-separated file. The input parameters used in this analysis are the same as Table S4 except for the host name – ‘homo_sapiens’ and genome assembly version– ‘GrCh38’. Summary statistics on the editability are included as Table S7 and gRNA libraries are included as Supplementary data 3.

## RESULTS

### Design of *beditor* workflow

The *beditor* workflow contains sequential steps that lead from input target sequences and other input parameters to the designed gRNA libraries as output (Figure 1a). The user provides information about the desired set of mutations as an input and a library of gRNAs is generated with the corresponding *a priori* estimates called *beditor* scores to help users in selecting the best performing gRNAs. *beditor* can also be used to execute only a subset of the analysis steps by changing the input parameters or providing inputs for intermediate steps. The standard input of *beditor* depends on the format of mutations i.e. nucleotide or amino acid. To carry out nucleotide level mutations, the users need to provide genome coordinates and the desired nucleotide after mutagenesis. For carrying out amino acid level mutations, the users provide Ensembl stable transcript ids, the position of the targeted residues and the corresponding mutated residue. Users can also provide inputs to limit the amino acid substitutions to a custom substitution matrix and specify whether only non-synonymous or synonymous substitutions should be carried out. In addition to creating mutations on a wild-type background (‘create’ mode), the *beditor* workflow also provides an option to design guides that would remove alternative SNPs and to mutate to the reference or wild-type alleles (‘remove’ mode). The program can be accessed via GUI (Figure S3), command line or API. In addition to the gRNAs designed to carry out provided mutations, the *beditor* workflow can also design control gRNAs that are important in the large-scale mutagenesis experiments. The positive control gRNAs designed by *beditor* generate non-sense mutations in the coding region of interest and negative control gRNAs that lack an editable nucleotide in the editing window of the BE, thus expected to have null effect.

### Customizability for broad utility

The *beditor* workflow utilizes a PyEnsembl python API (19) to fetch and work with the genomes of over 125 species and their various assemblies from the Ensembl genome database(18, 25), providing a broad utility for researchers across a wide spectrum of fields. *beditor* is also compatible with any custom user-made annotated genome. The ability to carry out parallel processing allows for the design of large gRNA sequence libraries using minimal computational resources (Figure S4). The users can incorporate BEs with varied editing properties and even novel BEs as per requirements. Similarly, they can incorporate any custom PAM sequences (for example Table S3) in addition to the experimentally validated PAMs already incorporated in the current version of *beditor* (listed in Table S8). Additionally, the location of the PAM with respect to the gRNA (upstream or downstream) and the provided length of gRNA are taken into consideration in the analysis. Lastly, *beditor* scores allows the users to select the best set of gRNAs for mutagenesis experiments.

### Selecting the best performing gRNAs

We defined a novel *beditor* scoring system that can be used to select the best performing gRNAs from designed gRNA library. Due to the lack of large scale, genome-wide base editing data, we relied on few general rules that are applicable to all BEs. These rules pertain to the requirements for optimal mutagenesis. With the penalties assigned to each requirement, the *beditor* score of the optimally performing guide RNA would tend to be higher while that of poorly performing guide RNA would be lower (see Methods). In the future, we wish to determine the values of handcrafted penalty scores from empirical data. The current version of *beditor* scoring system assesses four of the general requirements for optimal mutagenesis. Based on the conformity of the gRNA to the requirements four penalties scores are assigned (Figure 1b, see Methods). (1) While general rules of the BE mediated editing is an active field of research, we utilize the basic rule which is common between all base editors: the editable base should lie within the maximum activity window of the BE. Thereby, the gRNAs are penalized if the editable base does not lie within the maximum activity window of BE. Next, utilizing the alignments of gRNAs to the genome, potential off-target sites are identified. (2) A penalty score is assigned based on the gRNA binding at off-target. It is evaluated by capturing a general trend of mismatch tolerance along the length of a gRNA (24) (See Methods). Given the lack of large scale, base editing data, we used empirical data from ‘conventional’ CRISPR-Cas9– based genetic screens, assuming that the basic principles of gRNA recruitment and binding would be conserved between the two variants of CRISPR based mutagenesis technologies. (3) Additionally, from alignments of the gRNAs, a penalty is assigned based on the location of the off-target site (genic or intergenic regions). Off-target editing at intergenic regions is less likely to confound the mutational effects compared to functionally important genic regions. Accordingly, penalties are assigned. (4) Lastly, in order to account for number of off-target sites, the 3 penalties are multiplied together to evaluate a *beditor* score per gRNA. Effectively, the optimal gRNAs have a *beditor* score of 1, while a lower *beditor* score indicate incompatibility with BE requirements, significant off-targets or off-target effects that confound mutational effects. Finally, in order to filter out the gRNAs containing putative RNA polymerase III transcriptional terminators (26), the length of the poly-T stretch per individual gRNA is indicated in the output of *beditor*.

### Demonstrative analysis 1: customizability with respect to base editing strategies

We demonstrate that the users can incorporate and use custom base editors and PAM recognition sequences by designing gRNA libraries against sets of 1000 randomly assigned mutations (Supplementary methods) with 6 base editors (among which 4 are not yet discovered and are hence called “hypothetical”) and 16 PAM recognition sequences. With all the combinations of BE specific requirements and all combinations of two mutation formats (nucleotide and amino acid) and two modes of mutagenesis (‘remove’ and ‘create’), gRNA libraries were designed (Figure 2, and Supplementary data 1).

### Demonstrative analysis 2: ability to work with different species

To show *beditor*’s ability to work with genomes of different species, we designed gRNA libraries for representative 10 Ensembl genomes, for sets of 1000 randomly assigned mutations (Supplementary methods). The genomes of all the species were directly fetched from the Ensembl Genome database and gRNA libraries were designed for both two mutation formats (nucleotide and amino acid) and two modes of mutagenesis (‘remove’ and ‘create’). Through the designed gRNA libraries (Supplementary data 2) reasonable editability (Table S6) was achieved for all the species.

### Case study: designing a gRNA library against a set clinically relevant SNPs

To demonstrate the utility of our computational workflow, we designed a library of gRNAs against a set of clinically relevant SNPs in the human genome composed of 61,083 nucleotide level and 81,819 amino acid level mutations (see Supplementary methods). This analysis was carried out with two different BEs: Target-AID and ABE, and two PAM sequences: NGG (27, 28) and NG (29, 30) and in ‘create’ and ‘reverse’ mode. Note that the purpose of this case study analysis is to provide a demonstration of the functionalities of the *beditor* workflow. For instance, ABE base editor is not known to function in association with NG PAM. Yet, as shown in demonstrative analysis 1, the users may try any combinations of custom base editors and PAM sequences, The output libraries of gRNA sequences (Supplementary data 3) targets ~25% of the total mutations provided as input (Table S7). The resulting gRNA libraries were composed of gRNAs designed with each of the input BEs and PAM sequence which targeted both the strands (Figure 3 a and b). On average, ~1.6 guides were designed for each mutation. Summary visualizations of gRNA libraries (Figure 3 c) as well as visualizations of alignments of gRNAs with the target sequence (Figure S5) were generated. Additionally, the percentage of substitutions that can be edited with the designed guides (% editability) is represented as substitution maps (Figure 4) to indicate the proportion of input mutations that can be edited with the input BEs and PAMs. The gRNAs designed for this case study analysis as well as with all the combinations experimentally validated pairs of base editors and PAM sequences (Table S8) are provided as a database with a web interface at http://rraadd88.github.io/soft/beditor.

**Figure 3.**
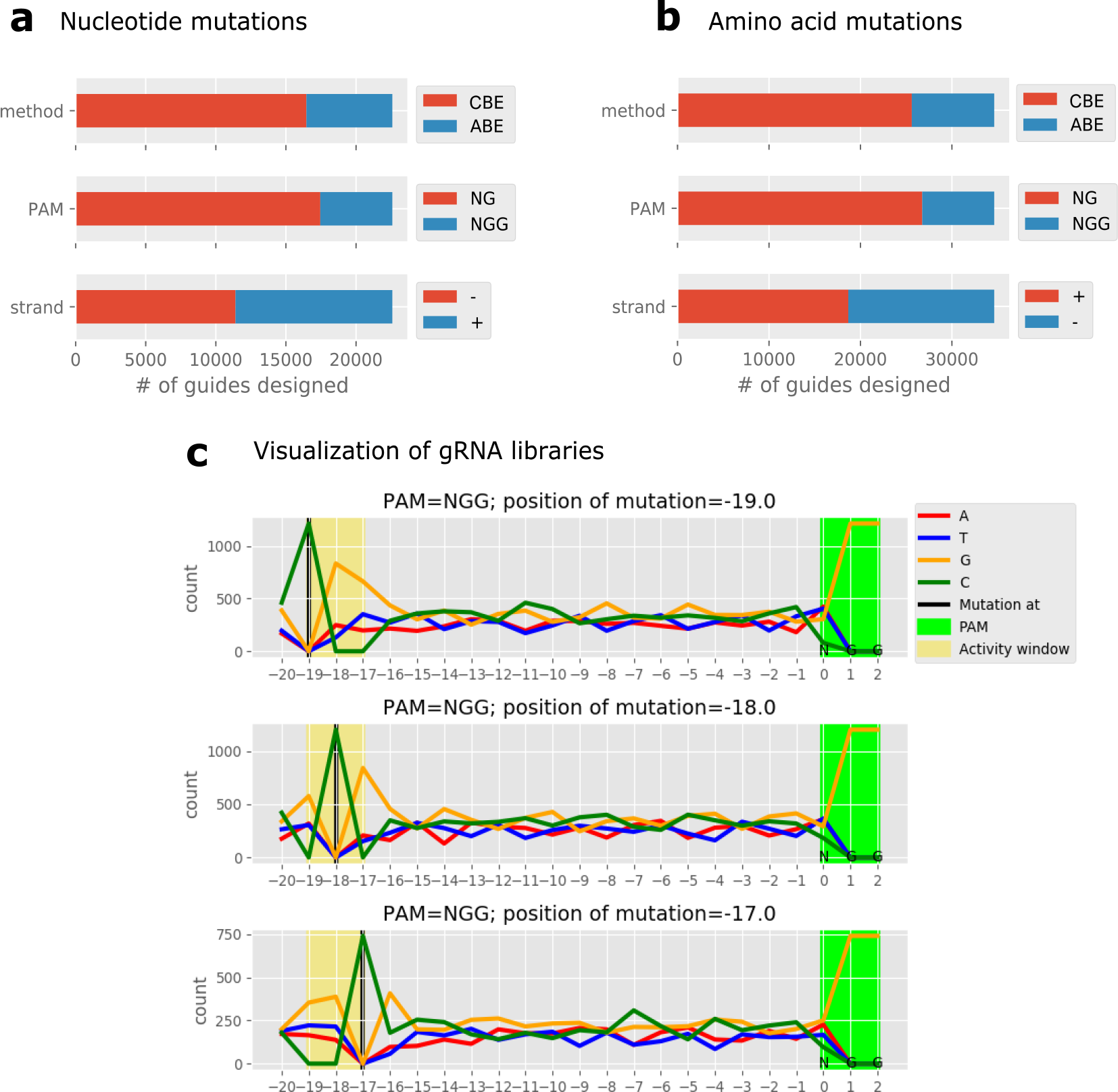
Case study analysis of clinically relevant human SNPs. For the case study analysis, 2 base-editors (Target-AID and ABE) and 2 PAM sequences (NGG and NG) were used. Number of gRNAs designed using each mutation format i.e. nucleotide (**a**) and amino acid mutation (**b**) data are shown. **c** Representative summary visualization of gRNA libraries designed with Target-AID base editor. Nucleotide composition of the gRNAs is shown along the length of the gRNAs. gRNAs are grouped by the position of the editable nucleotides within the activity window of a BE (shown in the rows).

**Figure 4.**
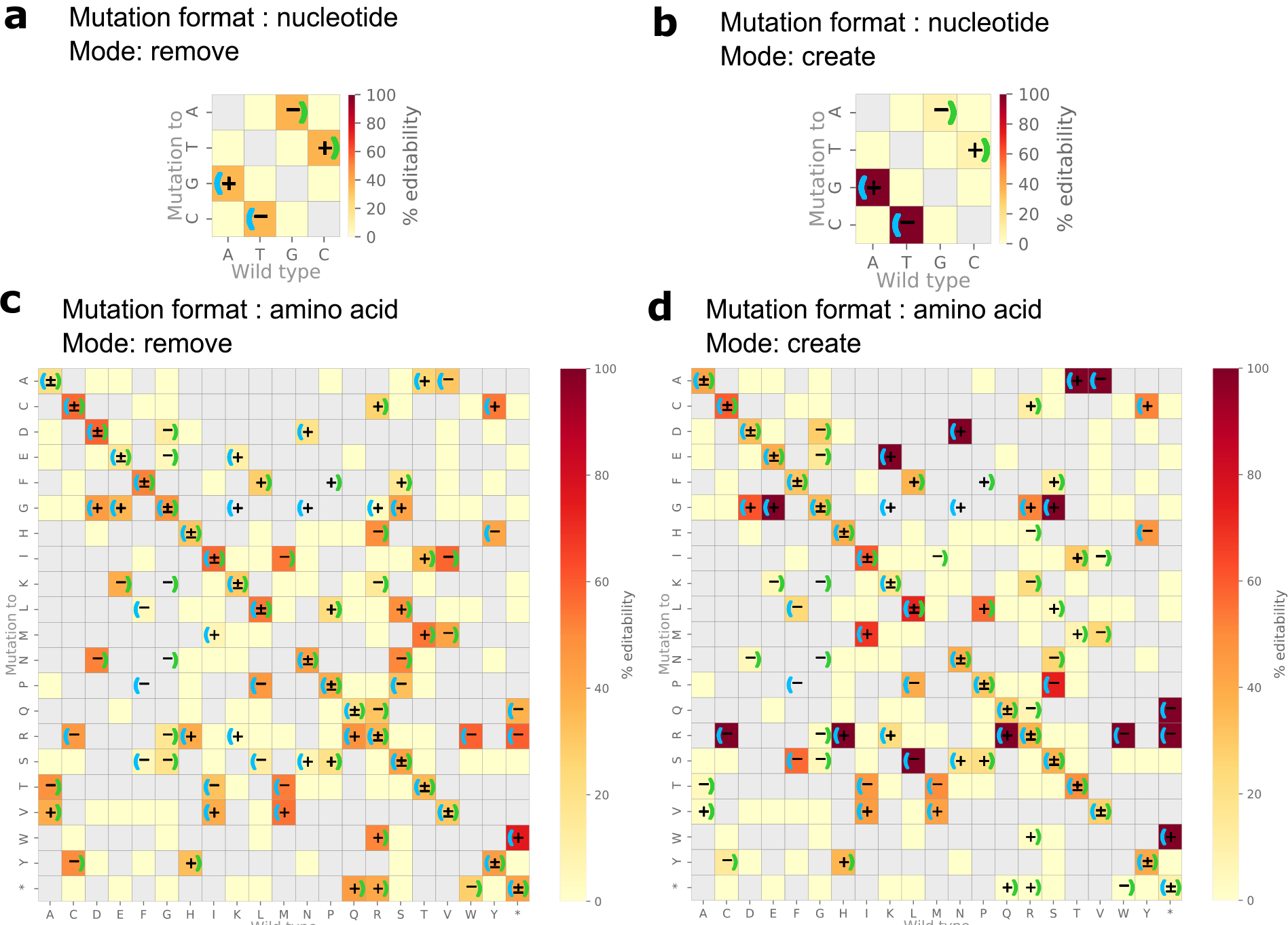
Percentage editability of gRNA libraries designed in case study analysis of clinically relevant human SNPs. Percentage of substitutions that can be edited by gRNA library (% editability) designed for case study analysis of clinically relevant human SNPs, in the format of nucleotide (**a** and **b**) and amino acid mutations (**c** and **d**) (see Supplementary methods). Also, the gRNA libraries were designed to remove mutations i.e. ‘remove’ mode (**a** and **c**) and to introduce mutations i.e. ‘create’ mode (**b** and **d**). Mapped on the heatmaps is a ratio between number of substitutions that can be edited with the designed gRNAs and the number of substitutions present in the input data (% editability). Left and right brackets indicate that the substitution is carried out by ABE and Target-AID respectively. +, − and ± indicate substitutions for which gRNA is designed on +, − and both strands respectively. Shown in gray are substitutions that are absent in the input data. * indicates non-sense mutation.

### Performance evaluation of *beditor* score

From the case study analysis, *beditor* scores were evaluated for each gRNA sequence in the library. From the distribution of scores (Figure S6), the users may assign a threshold to filter out gRNAs with lower *beditor* scores. Collectively, by definition, the *beditor* scores are negatively correlated (ρ=−0.94) with the number of off-target alignments (Figure 5a) and penalty assigned for each alignment based on distance of mismatches from the PAM sequence is positively correlated (ρ=0.65) with the distance (Figure 5b). Note that the rank correlation is not perfect because of cases in which there were two mutations in the aligned sequence. Also, purely informed from features of alignments of gRNAs and the requirements of BEs, the *beditor* score recapitulates empirical activity values of gRNAs with a strong positive correlation (ρ=0.95) determined in terms of Cutting Frequency Determination (CFD) score (24) (Figure 5c).

**Figure 5.**
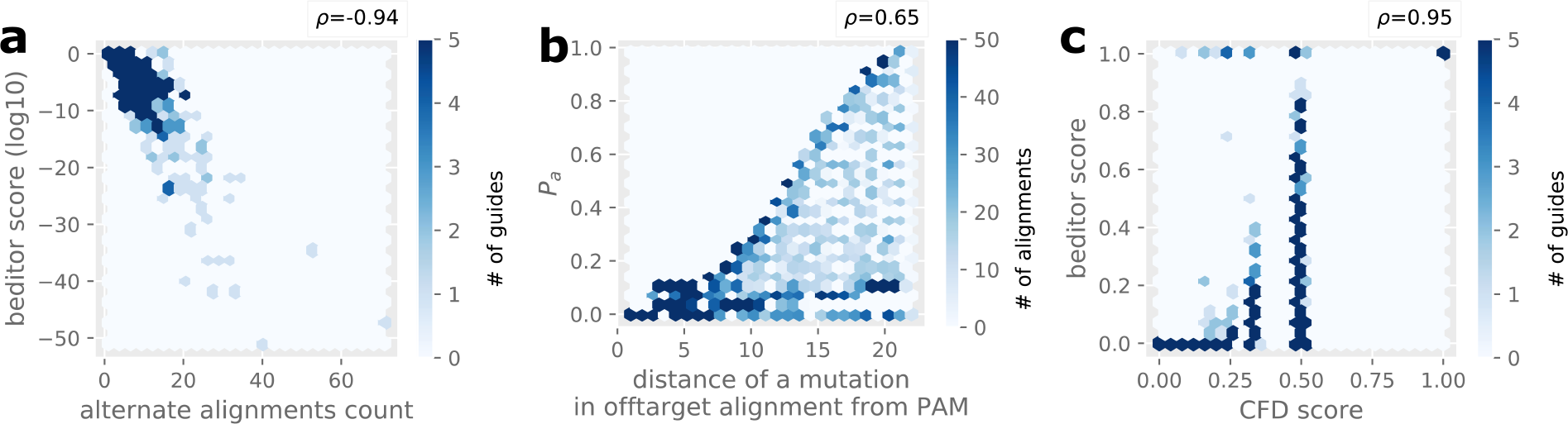
Performance assessment of beditor scores from case study analysis of clinically relevant human SNPs. **a** Relationship between the number of genome-wide off-target alignments and *beditor* score per gRNA. The color of hexbins are scaled according to the number of gRNAs per bin. **b** Relationship between the distance of a mutation in off-target alignments and corresponding penalty assigned (*P*_*a*_). The color of hexbins are scaled according to the number of off-target alignments per bin. **c** Relationship between the CFD score and *beditor* score for all the gRNAs carrying NGG PAM sequence. The color of hexbins are scaled according to the number of gRNAs per bin. *ρ* is Spearman’s correlation coefficient.

## DISCUSSION

CRISPR-mediated base editors have recently become a new paradigm in genome editing, owing to their unique ability to carry out precise mutations without the need for DNA breaks (5). Consequently, plethora of applications of BEs have emerged from all corners of the genome editing community, ranging from study of model and non-model organisms (3, 10–13) to therapeutics (14, 15). However, there has been a lack of robust and customizable software that can design gRNA libraries with any specific requirements of CRISPR-mediated base editing experiments. As presented here, the novel computational workflow of *beditor* (Figure 1a) fills in this important gap by allowing comprehensive customizability in terms of requirements of BE, PAM sequence and genome and thus increasing the applicability of the CRISPR-mediated base editing technology to broader community of researchers.

We demonstrate the modularity of *beditor* workflow in terms of the type of BE, PAM and genome by extensively testing them on synthetic sets of mutations. We also show that the workflow can be used to either create a mutation (‘create’ mode) or remove it (‘remove’ mode) with support for both nucleotide and amino acid format of mutations. Collectively, therefore, in terms of integrated customizability alone, *beditor* workflow provides a significant advance over other methods that provided only limited utilities (Table S1). In addition, we also introduce a novel method for *a priori* estimation of mutagenesis potential of gRNAs (Figure 1b), that utilizes features obtained from off-target alignments such as distance between mismatch and PAM recognition sequence. Such estimations would allow users to select a subset of designed gRNA library that would provide optimal mutagenesis in their experiments.

From the case study analysis of ~60,000 human clinically relevant SNPs, we show that *beditor* workflow provides all round gRNA design capabilities, scanning though combinations of multiple strategies (Figure 3a and b). The validations of *beditor* score from this analysis revealed that rather simple penalty-based evaluations efficiently captured dependence on position of the mismatch in the alignment from the PAM sequence (Figure 5b) and dependence of the off-target effects on number of alignments per gRNA (Figure 5a). Also, the estimations were in strong correlation with empirical data on the off-target effects obtained from ‘conventional’ CRISPR screens (Figure 5c). This is supported by the currently available data (31), that suggest that the Cas9 induced off target effects could be the predictors of off target effects of BEs (5). However, recent studies have found that the off-target effects of base editors can vary between ABE and CBE (32–34) and could even be largely unpredictable as in the case of CBEs containing APOBEC1 (33, 34). Therefore, in future, we wish to update the *beditor* scoring system further as more information would emerge from large-scale base editing experiments. The future developments of *beditor* workflow would be carried out in open source manner at https://github.com/rraadd88/beditor.

Together, considering the wide interest in scar-free and precise mutagenesis endowed by CRISPR-mediated base editing, moving ahead, novel and comprehensive gRNA designing workflow of *beditor* is expected to provide applicability to broad community of researchers and possibly become an essential component of the CRISPR-mediated base editing technology itself.

## DATA AVAILABILITY STATEMENT

The authors affirm that all data necessary for confirming the conclusions of this article are represented fully within the article and its tables and figures.

Processed data from this study i.e. gRNA libraries designed for demonstrative analysis 1, 2 and the case study analysis are provided as Supplementary data 1, 2 and 3 respectively. This dataset has been deposited using GSA Figshare portal.

The *beditor* software is available at https://github.com/rraadd88/beditor under the GNU General Public Licence (GPLv3). Database of gRNAs designed through *beditor* workflow can be accessed at http://rraadd88.github.io/soft/beditor.

The dataset analyzed in the study i.e. set of clinically associated human mutations were obtained from the Ensembl database in GVF format (ftp://ftp.ensembl.org/pub/release-93/variation/gvf/homo_sapiens/homo_sapiens_clinically_associated.gvf.gz, Date modified: 08/06/2018, 16:13:00).

## ACKNOWLEDGEMENT

We thank members of the Landry and Yachie Labs who provided important suggestions and feedback on the manuscript. We would like to thank Hideto Mori from the Yachie Lab for informative discussions.

## Supporting information

### Supporting figures

**Fig S1:**
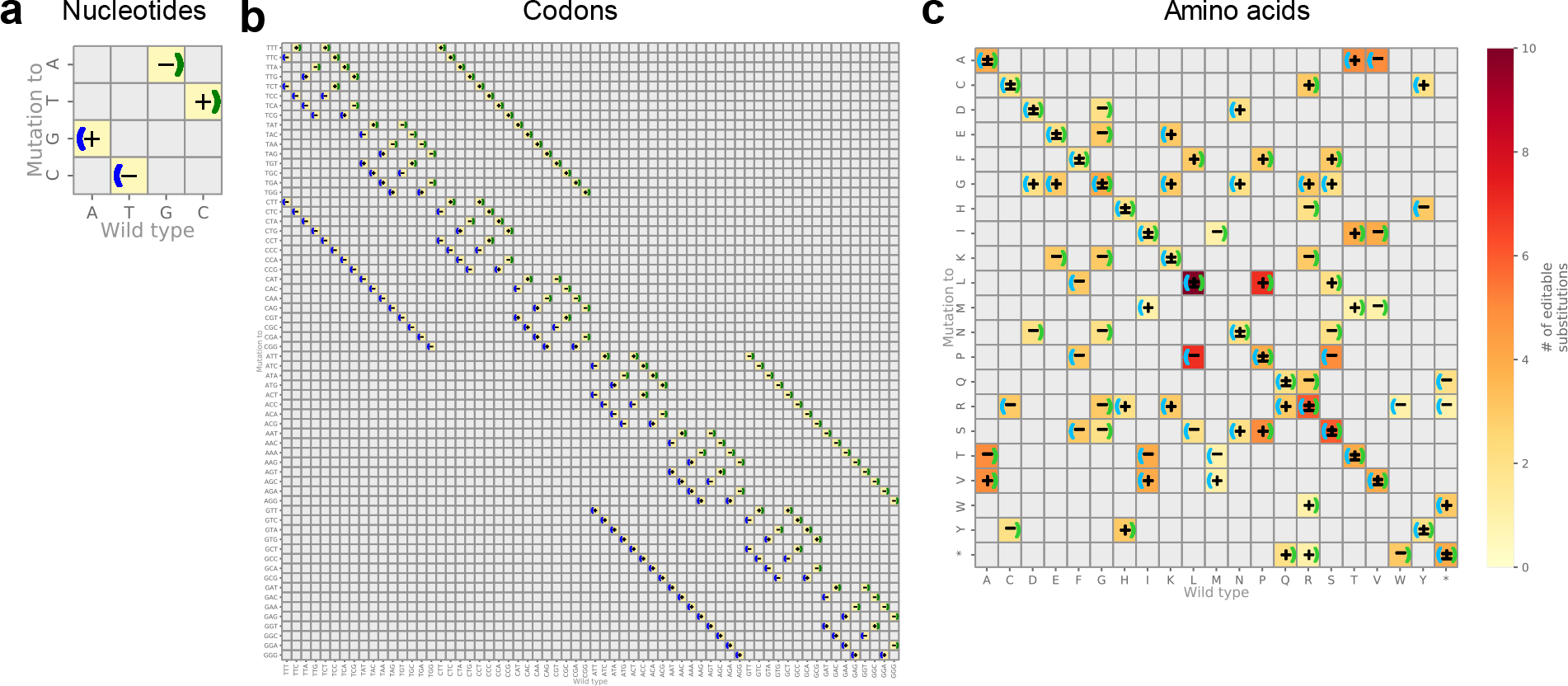
Possible substitutions by current ABE and CBE. **a** Nucleotide level substitutions. **b** Codon level substitutions. **b** Amino acid level substitutions. Shown on the heatmaps are the cumulative number of substitutions that can be edited with either ABE or CBE. Left and right brackets indicate that the substitution is carried out by ABE and CBE respectively. +, − and ± indicate substitutions for which guide RNA is designed on the +, − and both the strands respectively. Shown in gray are substitutions that are absent in the input data. The symbol * represents a non-sense mutation.

**Fig S2:**
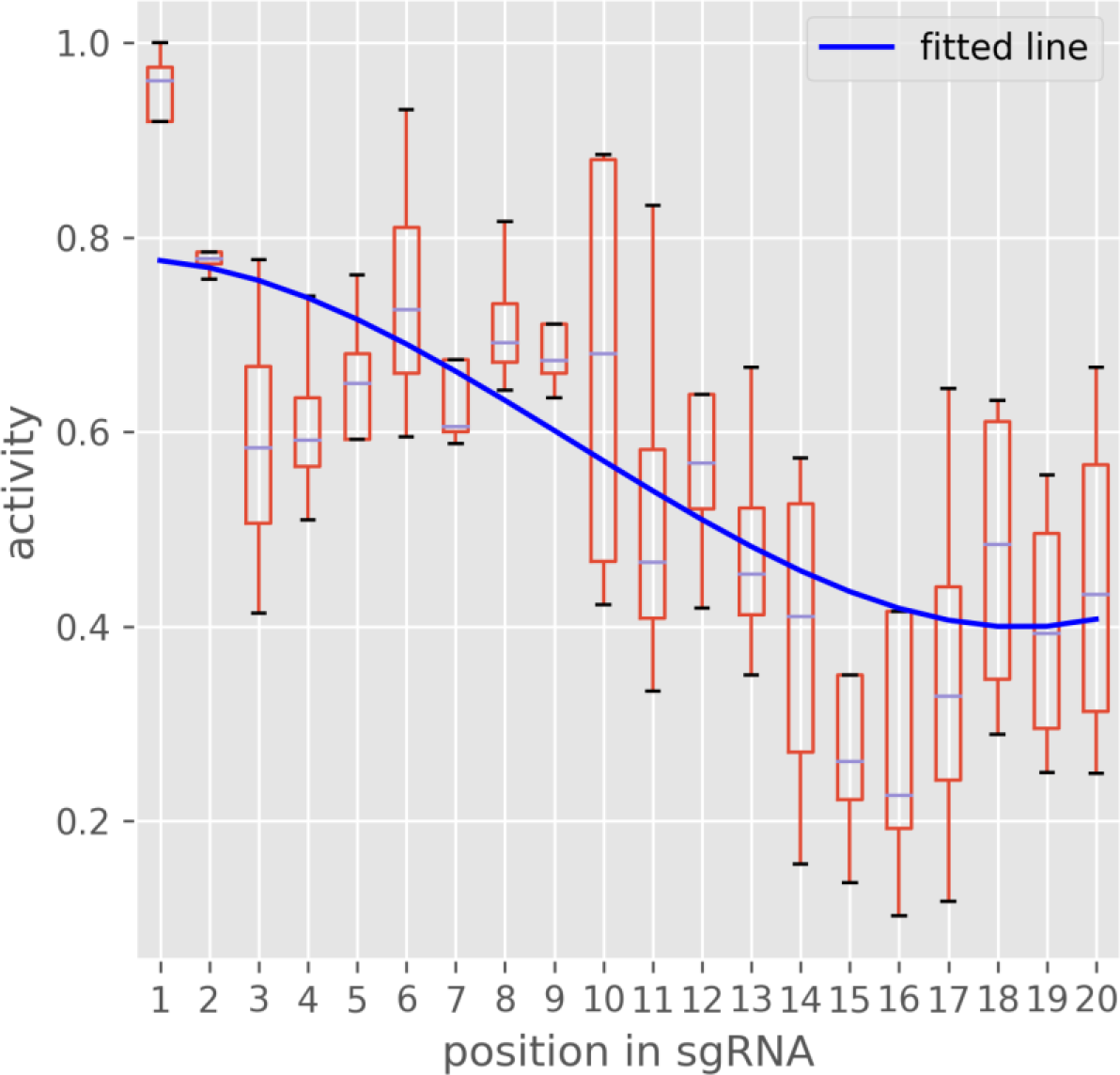
Fitting a third degree polynomial equation to the mismatch tolerance data from Doench et. al (3) The gRNA activity values at each mismatch position are shown as boxplots (red). The third degree polynomial equation fitted line is shown in blue.

**Fig S3:**
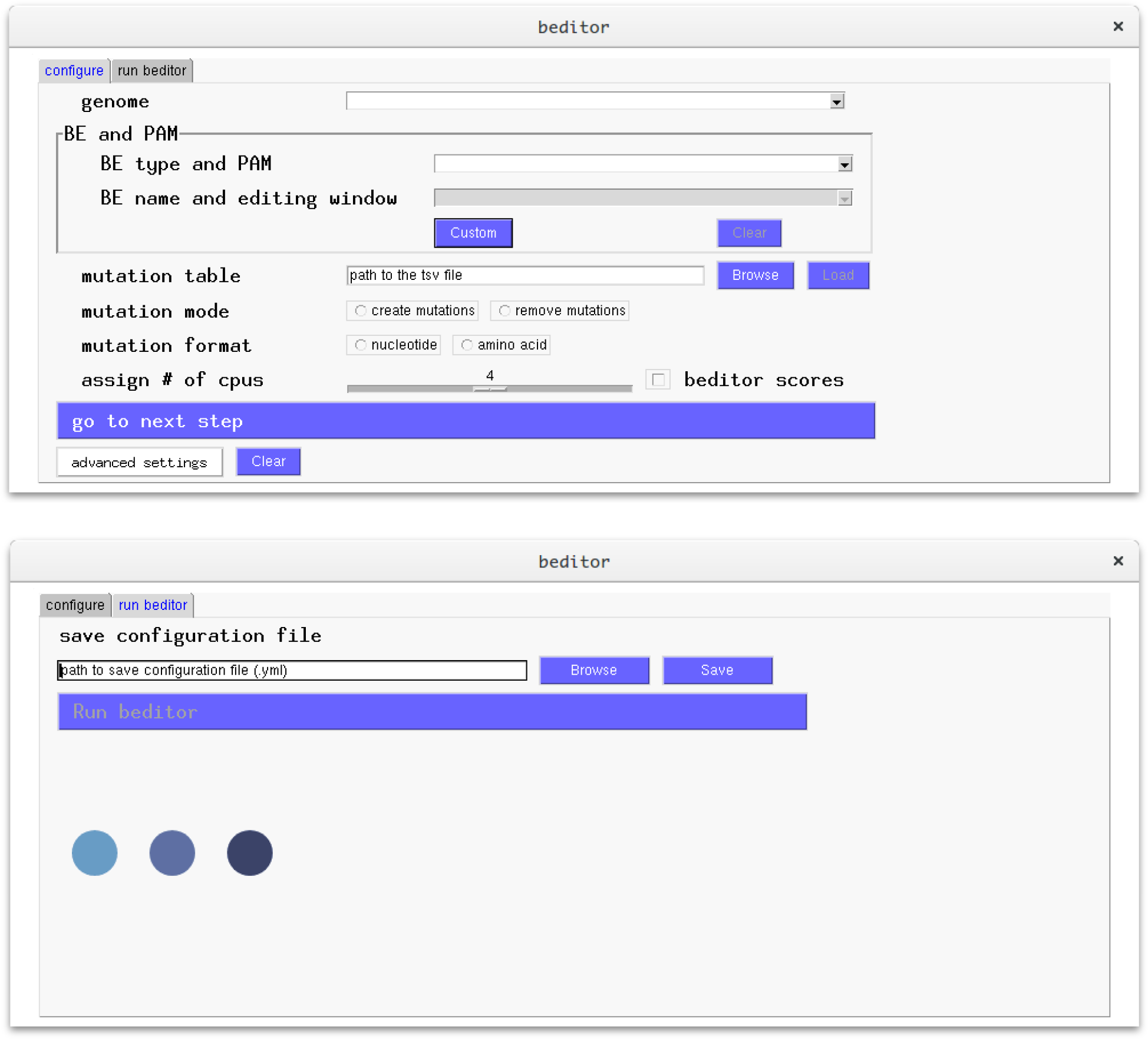
Graphical User Interface (GUI) of beditor. Top: In the first tab of the GUI, the parameters for the analysis workflow can be provided using a series of options. These basic parameters include the species name, the name of the base editor and the PAM sequence, an input list of mutations, the format of mutations (amino acids and nucleotide) and mode of mutagenesis (create or remove), the number of cores (processors) and If the *beditor* scores are to be calculated for the gRNAs. Bottom: the second tab of the GUI provides option to save the parameters of the analysis workflow as a YAML file and then subsequently run the program.

**Fig S4:**
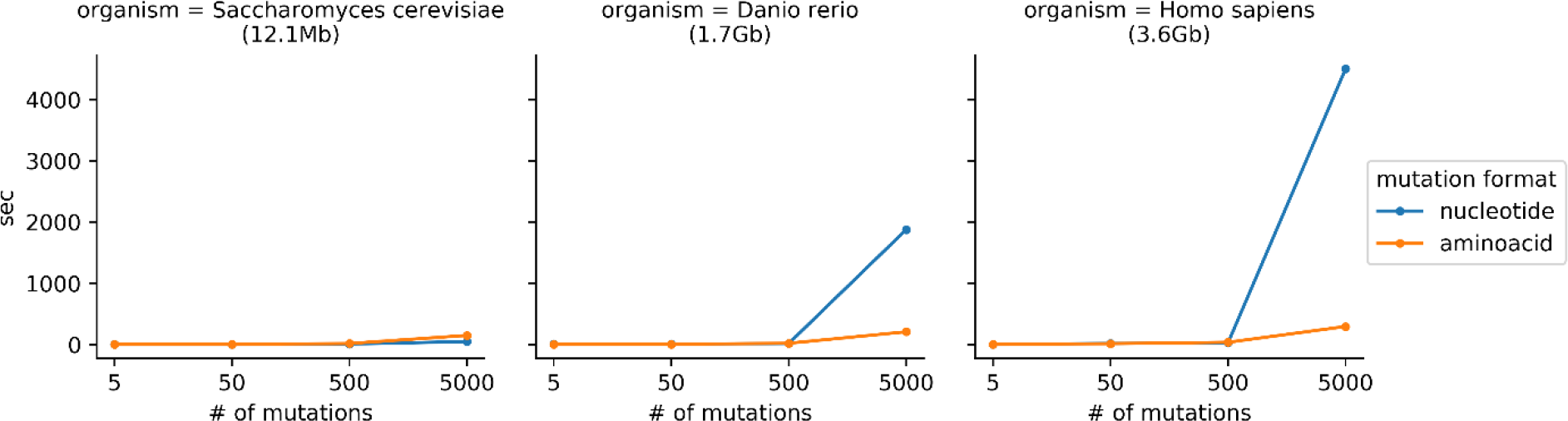
The time taken to design gRNA libraries depends on the number of mutations and the size of the genome. Sets of 5-5000 nucleotide and amino acid mutations were tested with genomes of 3 species. 6 parallel processors (cores) were used for the analysis. The analysis was carried out using test_beditor.py script from test_beditor repository (https://github.com/rraadd88/test_beditor). Note that the time does not include the time taken for installation of the genomes (i.e. downloading and indexing of the genomes).

**Fig S5:**
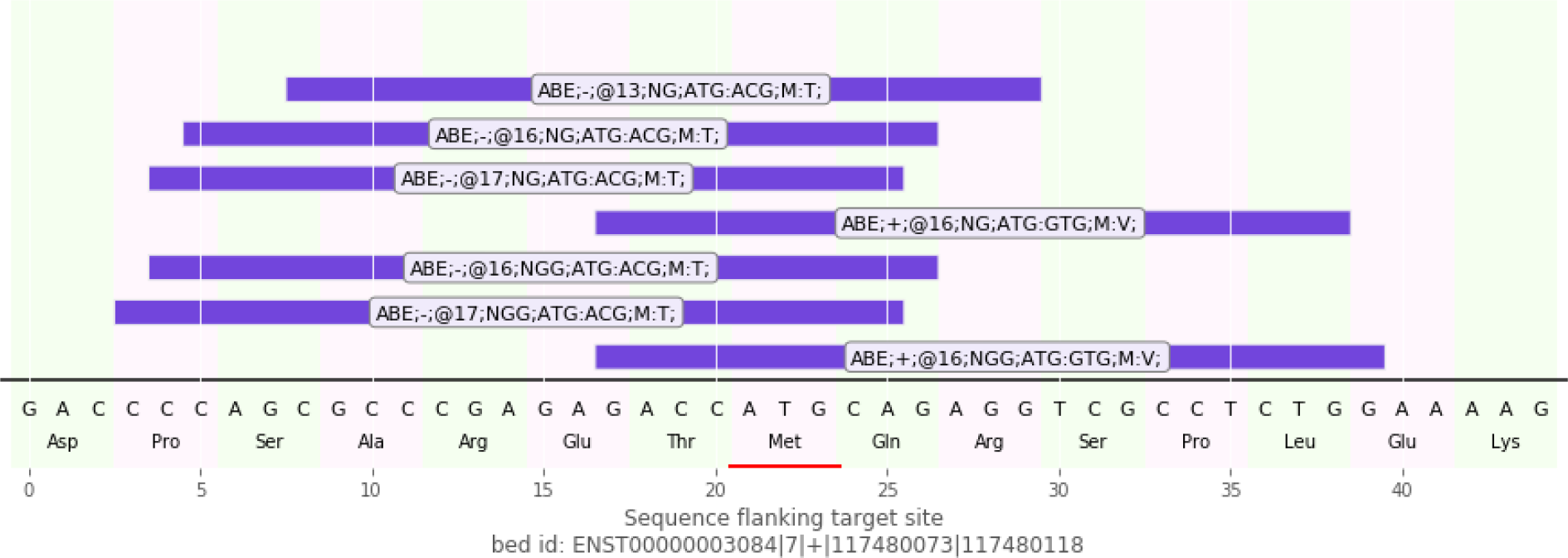
Representative visualizations of alignment between a gRNA and target genomic sequence. The gRNA are shown in purple and with annotations denoting its identity (name of the base editor, target strand, distance from PAM sequence, PAM, reference codon, mutated codon, wild-type amino acid and mutated amino acid) is shown on the guide RNA. The target site is indicated in red color. Reading frames and genomic coordinates of the target DNA are shown below.

**Fig S6:**
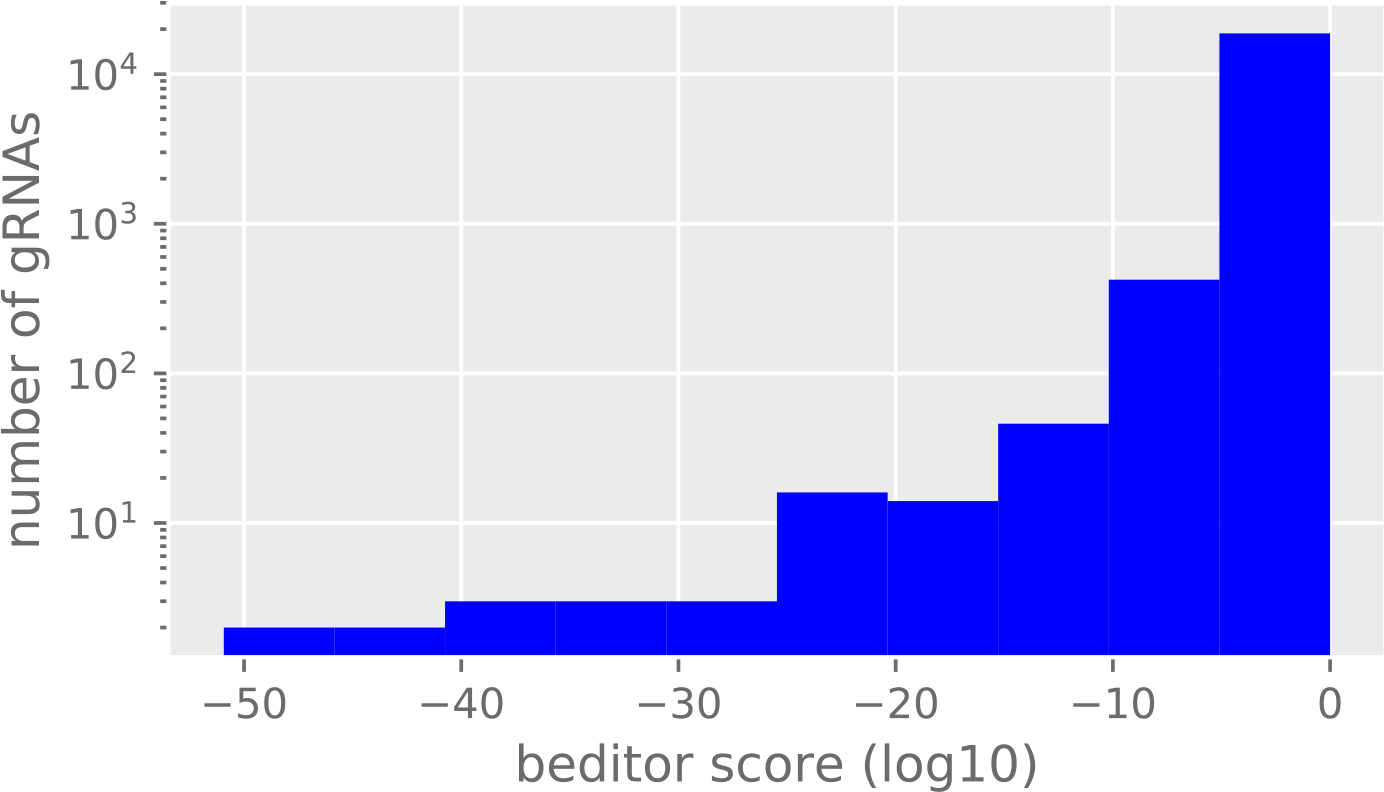
Distribution of beditor scores for the gRNA library designed for the demonstrative analysis of clinically associated human nucleotide mutations in ‘create’ mode.

### Supporting tables

**Table S1:** Comparison of features of *beditor* with other existing tools.

| Feature | *beditor* | iSTOP[1] | BE-analyzer[2] | Benchling [3] |
| --- | --- | --- | --- | --- |
| Number of species whose genomes are directly compatible | >125 | ~70 | ~40 | >100 |
| Searches sites editable by base editors | Yes | Yes | Yes | Yes only BE [4] |
| Designs gRNAs against predefined set of nucleotide mutations | Yes | No | No | No |
| Designs gRNAs against predefined set of amino acid mutations | Yes | No | No | No |
| Designs gRNAs for mutational scanning using a predefined amino acid substitution matrix | Yes | No | No | No |
| Designs gRNAs against non-sense mutations | Yes | Yes | No | Yes |
| Designs control gRNAs | Yes | No | No | No |
| Generation of gRNA libraries for genome-wide screening of mutations | Yes | No | 1 target site at a time | 1 target site at a time |
| Multithreading/ parallel computing | Yes | No | No | No |
| Allows addition of any novel/custom PAM recognition sequence | Yes | No | No | No |
| Allows addition of any novel/custom base editors | Yes | No | No | No |
| Estimation of editing efficiency of gRNAs | Yes | No | No | Yes |
| Integrates independent measures of off-target effects | Yes (CFD scores[5]) | No | No | Yes |

**Table S2:** Penalties used for the calculation of the *beditor* score.

| Penalty | Value |
| --- | --- |
| $P_{min}$ | 0.9 |
| $P_{max}$ | 0.1 |
| $G_g$ | 0.5 |
| $G_{ig}$ | 0.9 |
| $A$ | 0.5 |

**Table S3:** Default PAM recognition sequences supported by beditor.

| **PAM** | **Description** | **position** | **length of guide sequence** | **Reference** |
| --- | --- | --- | --- | --- |
| **NG** | xCas9 | 3' | 20 | [6] |
| **NGA** | SpCas9 mutant | 3' | 20 | [7] |
| **NGCG** | SpCas9 mutant | 3' | 20 | [7] |
| **NGG** | SpCas9 | 3' | 20 | [8,9] |
| **NGGNG** | Cas9 *S. Thermophilus* | 3' | 20 | [10,11] |
| **NGK** | xCas9 | 3' | 20 | [6] |
| **NGN** | xCas9 | 3' | 20 | [6] |
| **NNAGAA** | Cas9 *S. Thermophilus* | 3' | 20 | [10,11] |
| **NNGRRT** | SaCas9 | 3' | 21 | [12] |
| **NNNNACA** | CjCas9 | 3' | 20 | [13] |
| **NNNNGMTT** | Cas9 *N. Meningitidis* | 3' | 20 | [14] |
| **NNNRRT** | KKH SaCas9 | 3' | 21 | [7,12] |
| **TATV** | AsCpf1 mutant | 5' | 23 | [15] |
| **TTN** | Cpf1 *F. Novicida* | 5' | 23 | [16] |
| **TTTN** | Cpf1 *Acidaminococcus* / *Lachnospiraceae* | 5' | 23 | [16] |
| **TYCV** | TYCV AsCpf1 mutant | 5' | 23 | [15] |

**Table S4:**
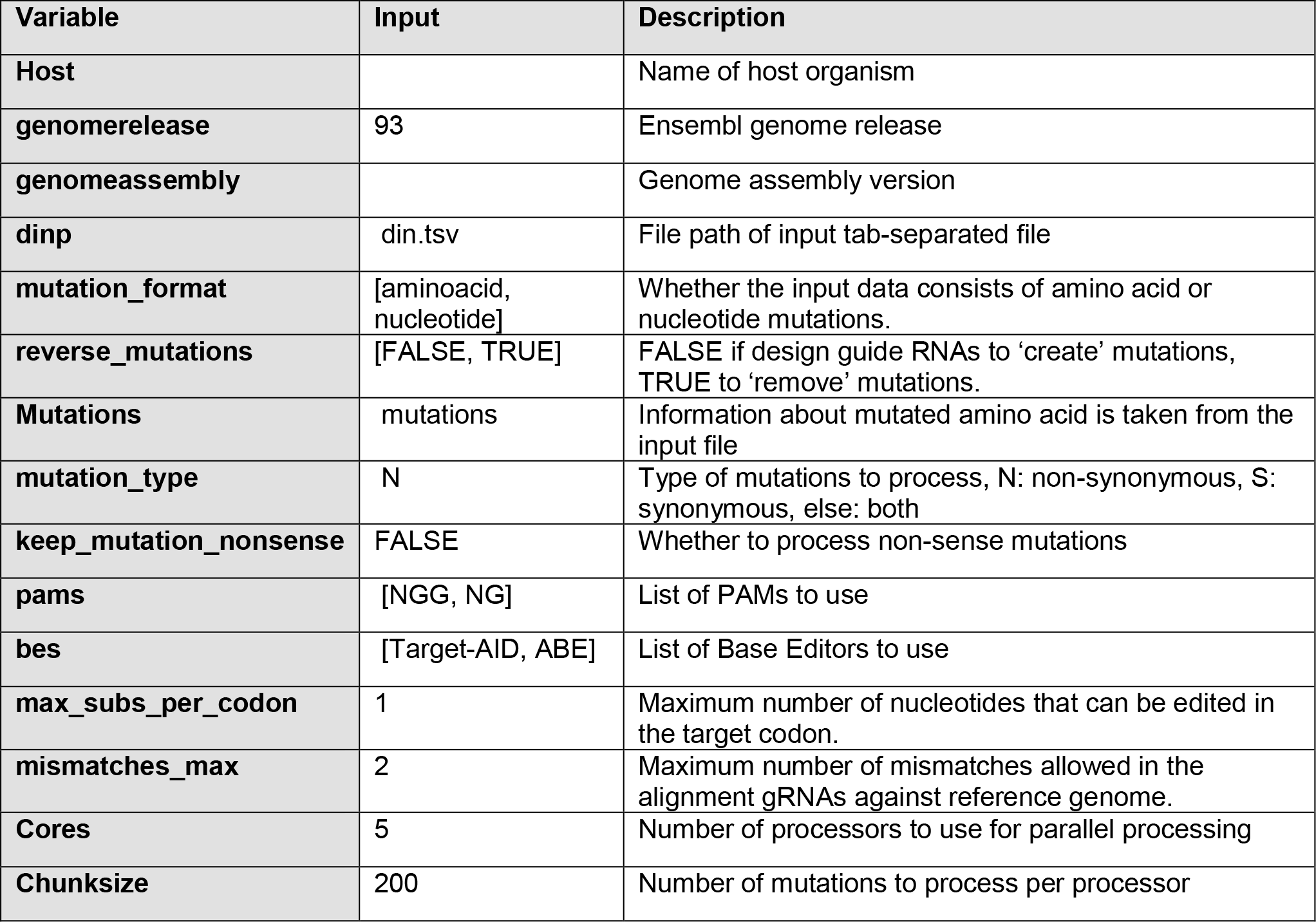
Input parameters used for the demonstrative analysis of clinically associated human genetic variants.

| Variable | Input | Description |
| --- | --- | --- |
| **Host** |  | Name of host organism |
| **genomerelease** | 93 | Ensembl genome release |
| **genomeassembly** |  | Genome assembly version |
| **dinp** | din.tsv | File path of input tab-separated file |
| **mutation_format** | [aminoacid, nucleotide] | Whether the input data consists of amino acid or nucleotide mutations. |
| **reverse_mutations** | [FALSE, TRUE] | FALSE if design guide RNAs to 'create' mutations, TRUE to 'remove' mutations. |
| **Mutations** | mutations | Information about mutated amino acid is taken from the input file |
| **mutation_type** | N | Type of mutations to process, N: non-synonymous, S: synonymous, else: both |
| **keep_mutation_nonsense** | FALSE | Whether to process non-sense mutations |
| **pams** | [NGG, NG] | List of PAMs to use |
| **bes** | [Target-AID, ABE] | List of Base Editors to use |
| **max_subs_per_codon** | 1 | Maximum number of nucleotides that can be edited in the target codon. |
| **mismatches_max** | 2 | Maximum number of mismatches allowed in the alignment gRNAs against reference genome. |
| **Cores** | 5 | Number of processors to use for parallel processing |
| **Chunksize** | 200 | Number of mutations to process per processor |

**Table S5:** Summary statistics of demonstrative analysis with custom base editors and PAM recognition sequences.

|  | nucleotide | amino acid | nucleotide | amino acid |
| --- | --- | --- | --- | --- |
| **Remove/create mutation** | remove | remove | create | create |
| **Total number of mutations in the input data** | 1000 | 1000 | 1000 | 1000 |
| **Total number of mutations edited** | 263 | 117 | 246 | 105 |
| **Total number of guides designed** | 817 | 562 | 791 | 403 |
| **% editability** | 26.3 | 11.7 | 24.6 | 10.5 |
| **number of guides per mutation** | 3.10646 | 4.80342 | 3.21545 | 3.8381 |

**Table S6:**
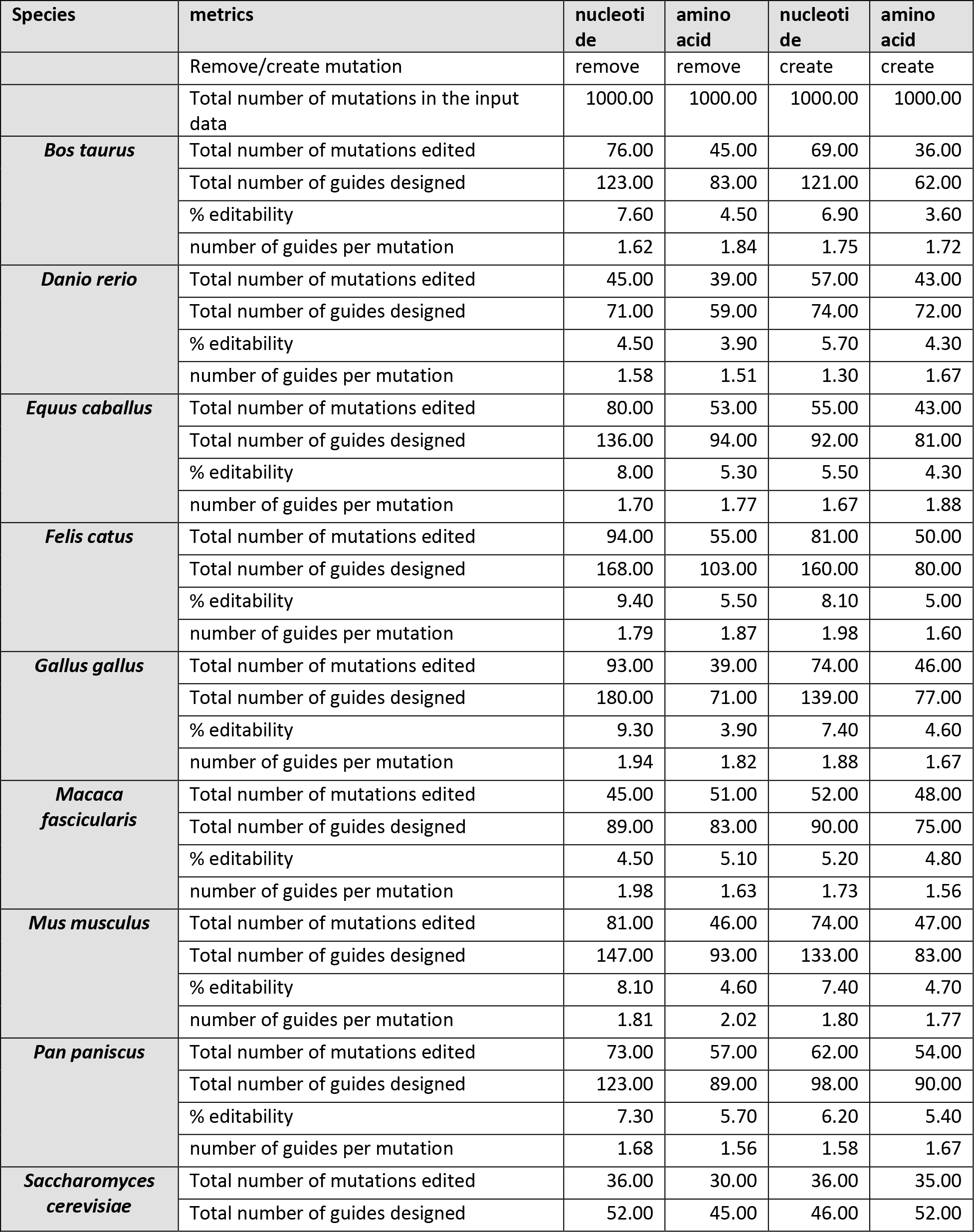
Summary statistics of demonstrative analysis of representative set of species.

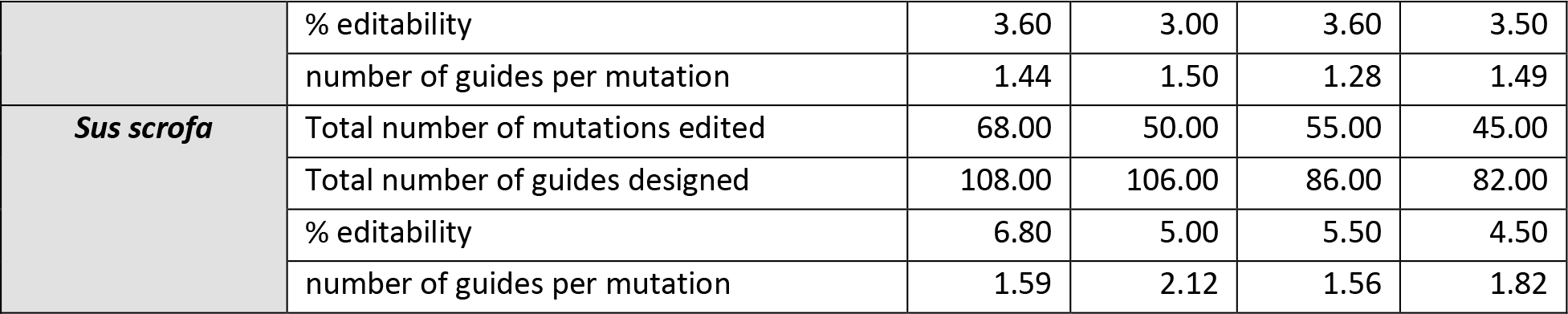

**Table S7:** Summary statistics for case study analysis of clinically relevant human SNPs.

| **Mutation format** | **nucleotide** | **amino acid** | **nucleotide** | **amino acid** |
| --- | --- | --- | --- | --- |
| **Remove/create mutation** | remove | remove | create | create |
| **Total number of mutations in the input data** | 61083 | 81819 | 61083 | 81819 |
| **Total number of mutations edited** | 13709 | 19996 | 13867 | 20420 |
| **Total number of guides designed** | 23432 | 35390 | 22587 | 33311 |
| **% editability** | 22.44 | 24.43 | 22.70 | 24.95 |

**Table S8:**
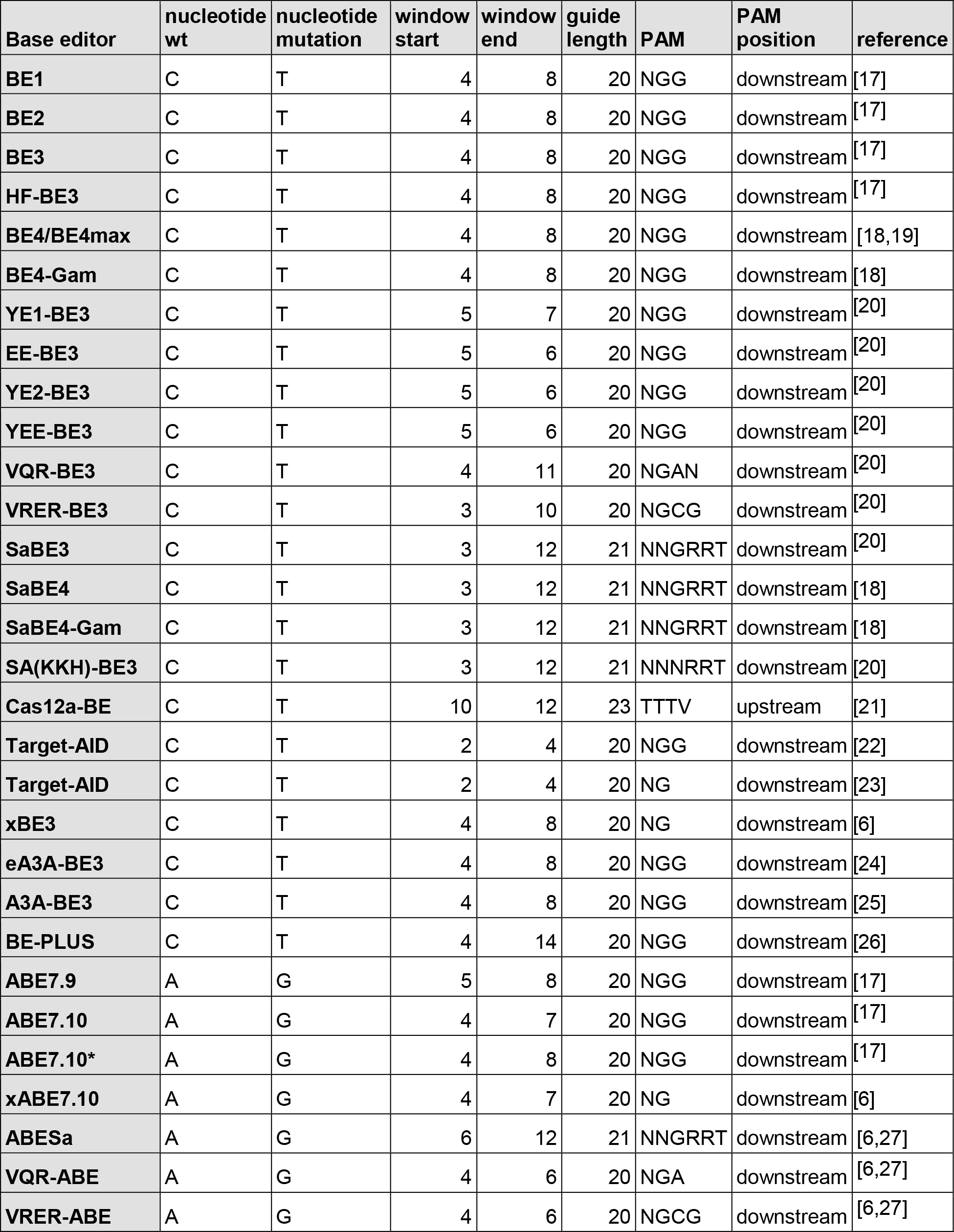
Default pairs of base editors and PAM recognition sequences supported by beditor.

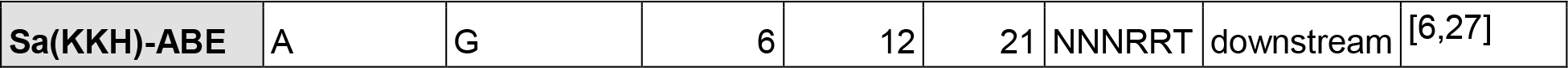

